# Prominent contributions of selective neuronal subtypes to pediatric drug-resistant focal dysplasia

**DOI:** 10.1101/2023.07.21.549980

**Authors:** Yujie Zhang, Lin Li, Fengjun Zhu, Xinyang Zhang, Dan Xia, Cheng Zhong, Shuli Liang, Dezhi Cao, Huafang Zou, Jing duan, Yousheng Shu, Yi Yao, Jianming Song, Songnian Hu, Jianxiang Liao, Qiang Zhou

**Author notes:** These authors contributed equally to this work (Yujie Zhang, Lin Li, Fengjun Zhu).

## Abstract

About 75% cases of epilepsy begin during childhood, and 10 - 30% of pediatric epilepsy patients are resistant to drug treatment. The predominant seizure type in children is focal cortical dysplasia (FCD) which is highly associated with drug-resistant epilepsy, but its underlying cause is poorly understood. We performed single-cell RNA sequencing and patch-clamp recording on fresh brain tissues obtained from pediatric FCD patients shortly after surgery to reveal critical factors contributing to FCD.

We report that known epilepsy genes or significant transcriptomic alterations occur in only a few neuronal subgroups, suggesting that epilepsy-associated neurons cluster into only a few neuronal subtypes. These epilepsy-subtypes displayed significant epilepsy-related transcriptomic alterations, especially in the genes associated with excitation/inhibition balance and neuron functional signal pathways. Among eight epilepsy-subtypes, L2_4_CUX2_YWHAH and PVALB_RGS5 subtypes showed the most prominent alterations in FCD patient tissues. Supporting PV-neurons being important, recording from fast-spiking/parvalbumin-containing neurons in acute FCD patient brain slices revealed reduced excitatory synaptic inputs, implicating lower activity in these neurons and fewer synaptic inputs onto them. A higher percentage of mitochondria genes, and likely higher activity in the pathways associated with oxidative phosphorylation and ATP biosynthetic process, was observed in the epilepsy-subtypes, suggesting a high energy expenditure in them. Interestingly, the activity of the above two pathways in most epilepsy-subtypes was lower in the FCD patients, suggesting these subtypes may be more vulnerable to energy deficit in epilepsy.

Altogether, transcriptomics is significantly altered in only a few neuronal subtypes in pediatric FCD patient brains. These selective epilepsy-subtypes may play prominent roles in the genesis of pediatric drug-resistant seizure and targeting them may provide new treatment options.

## Introduction

Epilepsy is the most common neurological illness and affects >1% of the world population.^1^ The prevalence and incidence of epilepsies in children is high (41 to 187/100,000), and about 75% of epilepsy cases begin during childhood.^2,3^ Drug-resistant epilepsy, which refers to the inability to fully control seizures with medication, affects approximately 1/3 of all patients with epilepsy, and 10 - 30% of children patients.^4,5^ Thus far, most studies have used adult epilepsy patient tissues, and much less are known about the alterations associated with epilepsy genesis in children. Since poorly controlled seizure severely affects the development and health of children, better treatments based on a deeper understanding of the cellular and molecular alterations in pediatric populations are of high importance.

Among newly diagnosed cases, focal epilepsy is the most prevalent form of epilepsy with an unknown etiology, with an annual incidence rate of 17.5/100,000 individuals.^6^ Focal cortical dysplasia (FCD) is one type of malformation of cortical development (MCD) characterized by disrupted cytoarchitecture of the cerebral cortex.^7^ FCDs are highly associated with drug-resistant epilepsy and are the predominant seizure type in children.^8,9^ More than 50% FCDs have seizure focus in the temporal lobe and are associated with temporal lobe epilepsy.^10,11^ In patients with epilepsy, about 61% have seizure onset prior to age 5, and 92.5% before age 16.^12^

Although FCDs and epilepsy have been linked since 1971, the pathophysiology of FCDs that causes epilepsy remains poorly understood.^13^ Imbalanced excitation and inhibition (E/I) ratio have been demonstrated in FCDs and is suggested to underlie epileptogenicity.^14,15^ This E/I imbalance can be caused by neurons with immature properties within the dysplastic tissue,^16^ and/or a lower density of GABAergic neurons.^17,18^ Mitochondria play a key role in supporting essential neurophysiological processes, including the maintenance of neuronal energy homeostasis and neuronal communication.^19^ The high activity of PV-neurons drives their high energy expenditure which heavily relies on mitochondrial energy metabolism.^20^ Mice with mitochondrial defects in PV-neurons display disinhibited circuits with imbalanced E/I.^21,22^ An increase in the extracellular ATP (eATP) levels in human hippocampal slices from drug-resistant epilepsy patients.^23^ or during spontaneous seizures, and a lower eATP level during the chronic phase, have been reported.^24^ Although these findings implicate the contribution of mitochondria/ATP signaling pathways in epilepsy, their exact roles and even directions of changes are poorly understood.

One interesting and potentially important recent finding is that the most prominent alterations in the epilepsy brains appear to occur in only a few selective neuronal subtypes. Both excitatory and inhibitory neurons can be divided into many subtypes. For example, a single functional human cortical area contains 45 inhibitory neuron and 24 excitatory neuron subtypes,^25,26^ while 13 principal (excitatory) neuron and 23 GABAergic (inhibitory) neuron transcriptomic subtypes were identified in the adult temporal cortex from temporal lobe epilepsy patients.^27^ In adult epilepsy, only a few families of principal neurons (L5-6_Fezf2 and L2-3_Cux2) and a small number of subtype somatostatin (SST) and parvalbumin (PV) neurons displayed significant transcriptomic alterations.^27^ This concept is potentially very important, as it suggests that epilepsy may be associated with or even caused by pathological alterations in only a few specific neuron subtypes. If this is true, focusing on these neuronal subtypes may provide crucial insights into both the genesis and treatment of epilepsy. Moreover, identifying the genes that are most significantly affected in these neuronal subtypes may provide effective treatments. Whether selective epilepsy-subtypes are also present in FCD or pediatric epilepsy patients is unknown.

Alterations in the GABAergic inhibitory neurons constitutes one major contributor to epilepsy.^28^ These inhibitory neurons include PV, SST and vasoactive intestinal peptide (VIP) positive neurons. In FCD patient epileptic tissues, a reduced expression of PV-immunofluorescence,^29,30^ an increased VIP neuron density but unaltered SST-neuron density^30^ were reported. Optogenetic inhibition of VIP-neurons disrupts seizure initiation and maintenance, while inhibiting PV- and SOM-neurons consistently reduced seizure duration but had mixed effects on seizure initiation.^31^ These findings also support the involvement of selective neuronal subtypes in epilepsy. However, whether selective subtypes of inhibitory neurons have a significant contribution to the pathogenesis in pediatric drug-resistant FCDs is also unknown.

To address the potential contributions of neuronal subtypes to pediatric FCD epilepsy, single-nucleus (snRNA-seq) is required. Due to the complexity of cortical neurons, the most effective and likely most accurate method to identify neuronal subtypes is to examine their transcriptomic markers using snRNA-seq. This approach preserves the heterogeneity of cells, and can thus be used to discover new or rare cell subtypes. The snRNA-seq analysis has revealed selective transcriptomic changes in certain neuronal subtypes in adult epileptic brain,^27^ and selective enrichment of MCD gene sets in subtypes of astrocyte and oligodendrocyte from pediatric MCD brain tissues.^32^ Although post-mortem tissue samples have been a major source for RNA sequencing,^33,34^ information obtained using fresh tissues is better at revealing the dynamic changes occurring in the brain during disease development and progression.^35,36^ For instance, when comparing post-mortem tissue to fresh human brain tissue, a specific decrease in the quantity of neuronal activity-dependent transcripts was observed.^35^ This advantage of using fresh brain tissue may be especially important for analyzing genes known to be affected by deteriorated cell health, such as mitochondria genes. Lastly, it is crucial to use tissues from pediatric patients to examine potential contributions of development-related genes to epilepsy genesis in children. In this study, we performed snRNA-seq on fresh brain tissues from pediatric drug-resistant FCD patients.

## Materials and methods

### Sample collection

Human brain samples were collected from temporal lobectomies of child patients undergoing surgery for FCD treatment in Shenzhen Children’s Hospital. Detailed clinical information related to the patients is summarized in Supplementary Table 1. All participants were enrolled from April 2021 to July 2022. This study was approved by the Institutional Ethics Committee, and all enrolled patients and their guardians have provided informed consent.

### Sample preparation

Pre-cooled PBSE was used to wash the brain tissue (PBS buffer containing 2 mM EGTA). Nuclei isolation was then performed with GEXSCOPE® Nucleus Separation Solution (Singleron Biotechnologies, Nanjing, China) according to the manufacturer’s product manual. Isolated nuclei were resuspended with PBSE to 10^6^ nuclei per 400 μl, filtered with a 40 μm cell strainer, and counted with Trypan blue. Nuclei enriched in PBSE were stained with DAPI (1:1,000) (TermoFisher Scientifc, D1306). Nuclei were defined as DAPI-positive singlets.

### Single nucleus RNA-sequencing library preparation

The concentration of single nucleus suspension was adjusted to 3-4 × 10^5^ nuclei/mL in PBS. Loading single nucleus suspension onto a microfluidic chip (GEXSCOPE® Single Nucleus RNA-seq Kit, Singleron Biotechnologies) and constructing the snRNA-seq libraries were performed according to the manufacturer’s instructions (Singleron Biotechnologies). An Illumina Nova seq 6000 instrument was used to Sequence the resulting snRNA-seq libraries with 150 bp paired end reads.

### Primary analysis of raw read data

Gene expression matrixes were processed by raw reads from snRNA-seq with CeleScope v1.9.0 pipeline (https://github.com/singleron-RD/CeleScope). In brief, to remove low quality reads, raw reads were first processed using CeleScope with Cutadapt v1.17 and then trim poly-A tail and adapter sequences.^37^ After extracting cell barcode and UMI, STAR v2.6.1a^38^ was used to map reads to the reference genome GRCh38 (ensembl version 92 annotation). Then we acquired UMI counts and gene counts of each cell with featureCounts v2.0.1 software.^39^ UMI counts and gene counts of each cell were used to generate expression matrix files for subsequent analysis.

### Quality control, dimension-reduction and clustering

Scanpy v1.8.2^40^ was used for quality control, dimensionality reduction and clustering under Python 3.7. For each sample dataset, expression matrix was filtered by the following criteria: (1) cells with gene count less than 200 or with top 2% gene count were excluded; (2) cells with top 2% UMI count were excluded; (3) cells with mitochondrial content > 20% were excluded; (4) genes expressed in less than 5 cells were excluded.

We obtained 109155 cells for the downstream analyses with an average of 2180 genes and 4700 UMIs per cell. The raw count matrix was normalized by total counts per cell and logarithmically transformed into normalized data matrix. Top 2000 variable genes were selected by setting flavor = ‘seurat’. Principal Component Analysis (PCA) was performed on the scaled variable gene matrix, and top 20 principal components were used for clustering and dimensional reduction. Uniform Manifold Approximation and Projection (UMAP) algorithm was applied to visualize cells in a two-dimensional space.

### Differentially expressed genes (DEGs) analysis

To identify DEGs, the scanpy.tl.rank_genes_groups() function based on Wilcoxon rank sum test with default parameters was used. Genes expressed in more than 10% of the cells in at least one of the two groups and with |fold change| ≥1. Adjusted p value (*p. adjust*) was calculated using benjamini-hochberg correction. *P. adjust* < 0.05 was used as the criterion for statistical significance.

### Cell type annotation

The cell type identity of each cluster was identified based on the expression of canonical markers found in the DEGs using SynEcoSys database. Heatmaps/dot plots and violin plots displaying the expression of markers used to identify each cell type were generated by Seurat v3.1.2 DoHeatmap/DotPlot/Vlnplot.

### Pathway enrichment analysis

To investigate the potential functions of DEGs, Gene Ontology (GO) and Kyoto Encyclopedia of Genes and Genomes (KEGG) analysis were performed using “clusterProfiler” R package 3.16.1.^41^ Pathways with *p. adjust* value < 0.05 were considered as significantly enriched. Gene Ontology gene sets including molecular function (MF), biological process (BP), and cellular component (CC) categories were used as references.

To investigate the expression of some pathways in different neuronal subtypes, these pathways were collected and used as functional gene sets for U Cell Gene set scoring. Query genes were ranked in order of their expression levels in individual cells and U Cell scores are based on the Mann-Whitney U statistic.^42^ Score ratio of U cell is defined as (Ep - Ctrl) * 100/Ctrl.

### Whole cell recording

Human brain tissues were rapidly removed and placed in the ice-cold sucrose-ACSF (in mM): 126 sucrose, 2 MgSO_4_, 2.5 KCl, 1.25 NaH_2_PO_4_, 26 NaHCO_3_, 10 D-glucose, 2 CaCl_2_ gassed with 95% O_2_ and 5% CO_2_. Slices of 350 μm were cut with a DTK-1000 tissue slicer (DTK, Japan) in the ice-cold sucrose-ACSF. Slices were transferred to a holding chamber with normal ACSF containing (in mM): 126 NaCl, 2.5 KCl, 1.25 NaH_2_PO_4_, 26 NaHCO_3_, 25 D-glucose, 2 CaCl_2_, and 2 MgSO_4_, and allowed to recover for 40 min at 34.5°C, then kept at room temperature until use. During recordings, slices were perfused with modified ACSF (modified from normal ACSF; in mM: 126 NaCl, 3.5 KCl, 1.25 NaH_2_PO_4_, 26 NaHCO_3_, 25 D-glucose, 1 CaCl_2_, and 1 MgSO_4_,). Recordings were conducted in slices on an Olympus microscope (BX51WI) with a 40X water-immersion differential interference contrast objective, at 35 - 36°C with oxygenated modified ACSF (4 - 5 ml/min). Resistance of the recording pipette was 4 - 8 MΩ. Recording pipettes were filled with K^+^ gluconate based intracellular solution contains (in mM):128 K^+^ -gluconate, 10 NaCl, 2 MgCl_2_, 10 Hepes, 0.5 EGTA, 4 Na_2_ATP, and 0.4 NaGTP.

Recordings were made from fast-spiking neurons in a depth of 50 - 100 μm from slice surface. HEKA EPC10 double patch clamp amplifier was used. Signals were collected at a sampling rate of 10 KHz and filtered at 2 KHz. Neurons with holding current > - 200 pA (at - 70 mV) were excluded from the data analysis. To record spontaneous excitatory post-synaptic currents (sEPSCs), somatic whole-cell voltage clamp recording (at −70 mV) was obtained from fast-spiking neurons in the slices. All neurons were recorded for at least 5 min to collect sEPSCs. Whole cell current clamp recording of evoked spikes was performed using a series of 500 ms depolarizing current steps with 4 s intervals, and every step with an increment of 20 pA (from 0 to 200 pA).

We used a few parameters to examine AP properties. The first AP evoked by rheobase was used to characterize AP shape. AP threshold was determined by the third derivative of the AP found over the AP rising phase. AP amplitude was calculated by measuring the absolute maximum amplitude from AP peak to −70 mV (holding Vm). Afterhyperpolarization (AHP) was measured using AHP peak and latency. AHP peak is the difference in membrane potentials between AP threshold and trough (lowest point of an AP), while AHP latency is the time difference between AP peak and trough. More than 400 sEPSC events in each neuron were identified and analyzed using Mini Analysis software. Mean amplitude of sEPSC in each neuron was used for group analysis.

Fast-spiking neurons infused with calcein (4 mmol/L). Spiking pattern was used to identify the fast-spiking neurons, and sEPSCs were recorded at −70 mV. Recording pipettes were slowly removed, and a microscope was used to determine that the cell had been infused with green dye and remained on the brain slice.

To confirm that the recorded neurons is PV-positive, we performed immunofluorescence staining.^43^ Slices were fixed with 4 % paraformaldehyde for 24 hr, then were dehydrated in 30% sucrose for 48 hr at 4 ℃. The sections were washed with PBS three times and 10 mins each. These sections were blocked with 10% normal goat serum that contained 0.5% triton-100 in PBS for 1 hr at room temperature. Sections were incubated with rabbit anti-parvalbumin antibody (1:2000, Abcam) at 4℃ overnight. Sections were then washed three times in PBS and incubated with an Alexa Fluor 546 conjugated secondary antibodies (goat anti-rabbit 546; 1:400; Invitrogen) for 1 hr at room temperature. Sections were then washed in PBS, cover-slipped, and imaged on a confocal microscope (Nikon A1R).

### Statistics and repeatability

Unpaired Two-tailed Wilcoxon rank sum test was used for comparison of gene expression or gene signature between two groups. All statistical analyses and presentation were performed using R. *P* < 0.05 was considered as statistically significant.

### Data availability

The data supporting the findings of this study are available from the corresponding author on reasonable request.

## Results

### Distinct neuronal subtypes in brain tissues from pediatric FCD patients

The snRNA-seq was used to analyze tissues from the temporal lobe of pediatric FCD patients obtained during surgery (Fig. 1A). The epileptic focus zone was identified prior to surgery, and control tissues were from the paraepileptic zone removed to ensure adequate seizure control.^44^ We collected 10 samples from 10 patients in the epilepsy group and 7 in the control group (Supplementary Fig. 1A, 1B), matched for age and median genes (Supplementary Table 1, Supplementary Fig. 1C, 1D). We obtained a total of 103,345 single-nuclei gene expression profiles which have passed quality control (Supplementary Table 2).

**Figure 1.**
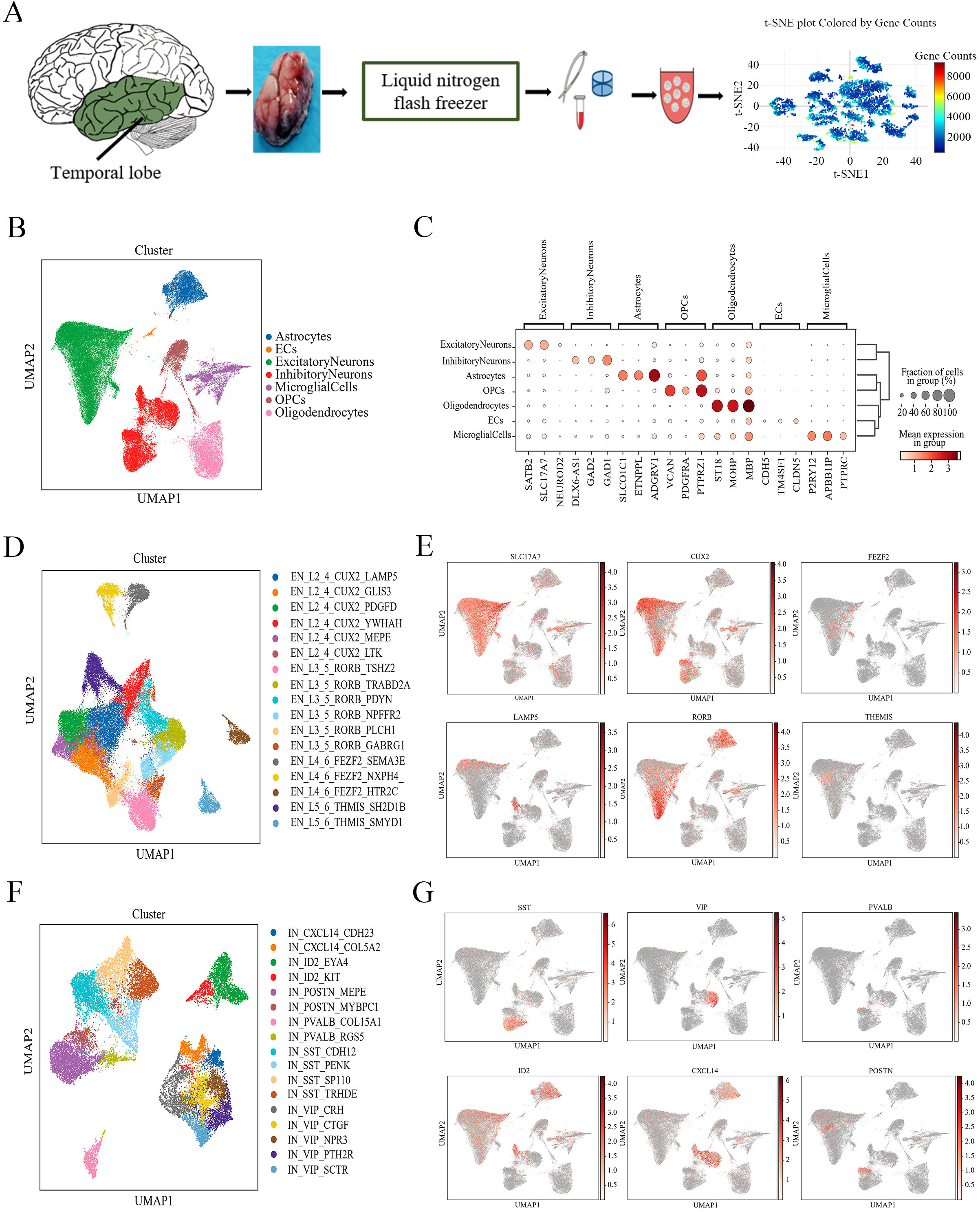
Overview of the single nucleus transcriptomic dataset. **(A)** Experimental procedure for snRNA-seq using brain tissue from child FCD patients. **(B)** UMAP representation of snRNA-seq data and cell type annotations based on the expression of known marker genes. **(C)** Expression of known marker genes for all cell types. **(D)** UMAP embedding and cell type annotation visualized for excitatory neuron subtypes. **(E)** Expression of general and family-specific marker genes in excitatory neurons. **(F)** UMAP embedding and cell type annotation visualized for inhibitory neuron subtypes. **(G)** Expression of general and family-specific marker genes in inhibitory neurons. For E and G, the redness of color is proportional to the log values of normalized expression level.

Seven distinct cell types were identified based on the expression of known cell type markers (Supplementary Table 3, Fig. 1B, 1C), with a total of 46,502 excitatory and 21,526 inhibitory neurons and each are further divided into 17 subtypes (Fig. 1D, 1F). Excitatory neurons were classified based on previously described layer-specific marker genes *CUX2* (L2_4)*, RORB* (L3_5)*, FEZF2*(L4_6), and *THEMIS* (L5_6), and inhibitory neurons were classified using cardinal marker genes *PVALB, SST, VIP, CXCL14* and *ID2* (Fig. 1E, 1G).^27^ We thus annotated 17 excitatory neuron and 17 inhibitory neuron subtypes. A third hierarchical level (for example, *YWHAH, PDYN*) was selected using the highest expressed DEGs between a given subtype and the rest (Fig. 1E, 1G, Supplementary Table 3).

### Selective enrichment of epilepsy genes in distinct neuronal subtypes

To examine whether significant expression of epilepsy-related genes occurs mostly in selective neuronal subtypes, we analyzed epilepsy gene set (C10012544) by comparing scores between pediatric FCD and control groups using U cell method (Fig. 2A, Supplementary Fig. 2A). Significantly higher scores were found in many neuronal subtypes in the FCD group (Fig. 2A, Supplementary Fig. 2A). We also obtained the same result using another epilepsy-related gene set from Wang *et al*.^45^ and Macnee *et al.*^46^ (Supplementary Fig. 2B, Supplementary Table 4-1). Closer inspection revealed that certain neuronal subtypes (L2_4_CUX2_LAMP5, L3_5_RORB_GABRG1, L3_5_RORB_PDYN, ID2_KIT, PVALB_RGS5, VIP_CRH and SST_PENK) have much higher score ratio of the FCD over control group (score ratio > 1, Fig. 2B, Supplementary Table 4-2), suggesting that these few subtypes may have more prominent contributions in epilepsy than the others. To test this hypothesis, we analyzed epilepsy genes that are associated with known epilepsy-causing mutations (Supplementary Table 4-3).^45^ While expression differences were detected in all neuronal subtypes, we discovered that the seven aforementioned subtypes, along with the L2_4_CUX2_YWHAH subtype, exhibited a significantly greater number of epilepsy genes with higher altered expression in the FCD group (number > 30, with avg_log_2_ (FC) > 0.25 or < −0.25) (Supplementary Table 4-3, Fig. 2C). Examination of the epilepsy-related genes revealed the same eight neuronal subtypes (Supplementary Fig. 2C, Supplementary Table 4-4).^46^ In addition, we identified genes with the largest altered expression levels (avg_log_2_ (FC) > 16 or < −16) between pediatric FCD and control groups, including *PNPO, TBC1D24, SLC13A5, CPA6, CASR, ADRA2B, GAL, LMNB2, NHLRC1, PRDM8, CHRNA2, DEPDC5* (Fig. 2D). Besides *LMNB2, NHLRC1, PRDM8, CHRNA2, DEPDC5*, *ADRA2B*, all other genes were selectively related to child epilepsy. Some genes showed altered expression in only one or several neuronal subtypes, with *PRDM8* showed higher expression in the CXCL14_COL5A2 subtype, *LMNB2* in ID2_KIT subtype, and *CASR* in all VIP subtypes (Fig. 2D).

**Figure 2.**
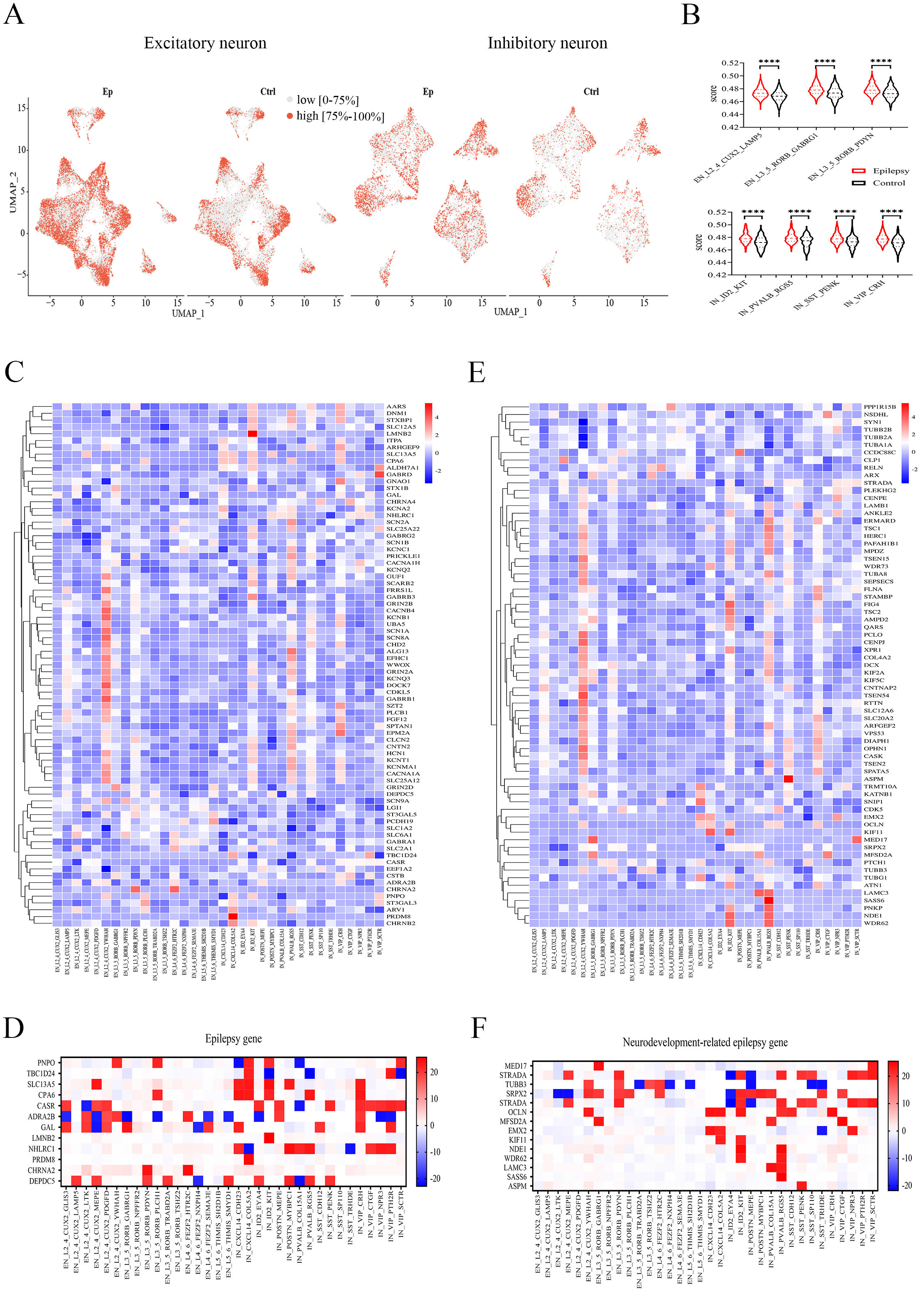
Epilepsy-related neuronal subtypes in FCD. **(A)** UMAP representation of the sores of epilepsy genes identified in DisGeNET (C10012544) in neuronal subtypes between epilepsy and control groups. Red represents high scores (top 75%) and gray represents low sores (0 - 75%). **(B)** Certain neuronal subtypes with high scores ratio between epilepsy and control groups. **(C)** Relative expression level (Ep vs. Ctrl) of epilepsy genes in excitatory or inhibitory neuronal subtypes. **(D)** Fold difference for significantly altered epilepsy genes between epilepsy and control groups. **(E)** Relative expression level (Ep vs. Ctrl) of neurodevelopment-associated epilepsy genes in excitatory or inhibitory neuronal subtypes. **(F)** Fold difference for significantly altered neurodevelopment-associated epilepsy genes between epilepsy and control groups. For B and D, the axis represents epilepsy or neurodevelopment-associated epilepsy genes, and the columns represent neuronal subtypes. Colors represent Z-score (average of log_2_ (Fold-Change)). Fold-Change = G1/G2, where G1 and G2 represent gene expression levels in epilepsy and control groups, respectively. For C and E, Colors represent average of log_2_ (Fold-Change).

The onset of epilepsy mainly occurs in childhood of FCD patients, with about 92% before age 16.^12^ The age range of our samples was between 5 months and 15 years, implicating a prominent role of neural development. We thus focused on genes with mutations known to be associated with neurodevelopmental malformations and epilepsy.^45^ In epilepsy patient tissues, we observed higher expression of many neurodevelopment-associated epilepsy genes that fall into the same eight neuronal subtypes identified above (Fig. 2E, 2F, Supplementary Table 4-5). Genes with significant difference between epilepsy and control groups (avg_log_2_ (FC) > 16 or < −16), including *OCLN, SRPX2, TUBB3, LAMC3, EMX2, NDE1, MED17, ASPM, MFSD2A, STRADA, KIF11, WDR62, S*ASS6, were identified (Fig. 2F). Among them, selectively higher expression was observed for *SASS6* in PVALB_RGS5 subtype and *LAMC3* in both PV-neuron subtypes (Fig. 2F). Interestingly, in both epilepsy gene set and neurodevelopment-associated epilepsy gene set, L2_4_CUX2_YWHAH and PVALB_RGS5 had the highest number of significantly altered genes (avg_log_2_ (FC) > 1 or < −1, *p* < 0.05). The above results collectively suggest that these eight subtypes are likely to have the most prominent contribution to the pathogenesis/pathology of epilepsy, and we thus termed them epilepsy-subtypes. Altogether, we have identified a few selective neuronal subtypes showing significantly higher altered expression of known epilepsy-related genes in the FCD patient brain tissues. This is the first report of epilepsy-subtypes in pediatric drug-resistant FCD epilepsy.

### Epilepsy-related signaling pathways and novel epilepsy-related genes in FCD tissues

What are the key functions of these significantly altered genes in the identified epilepsy-subtypes? Gene Ontology (GO) enrichment analysis on the DEGs in each subtype revealed that the most affected pathways are related to neuronal functions (Supplementary Fig. 3,). In the epilepsy-subtypes, more than 20 significantly enriched neuronal function-related GO terms have been identified. Higher than 60 GO terms associated with neuronal functions were observed in L2_4_CUX2_LAMP5, L2_4_CUX2_YWHAH, ID2_KIT, and PVALB_RGS5 subtypes (Fig. 3A). However, POSTN_MYBPC1 was not an epilepsy-subtype although having 59 significant GO terms. Thus, enrichment of neuronal function-related pathways alone is not sufficient to qualify as an epilepsy subtype.

**Figure 3.**
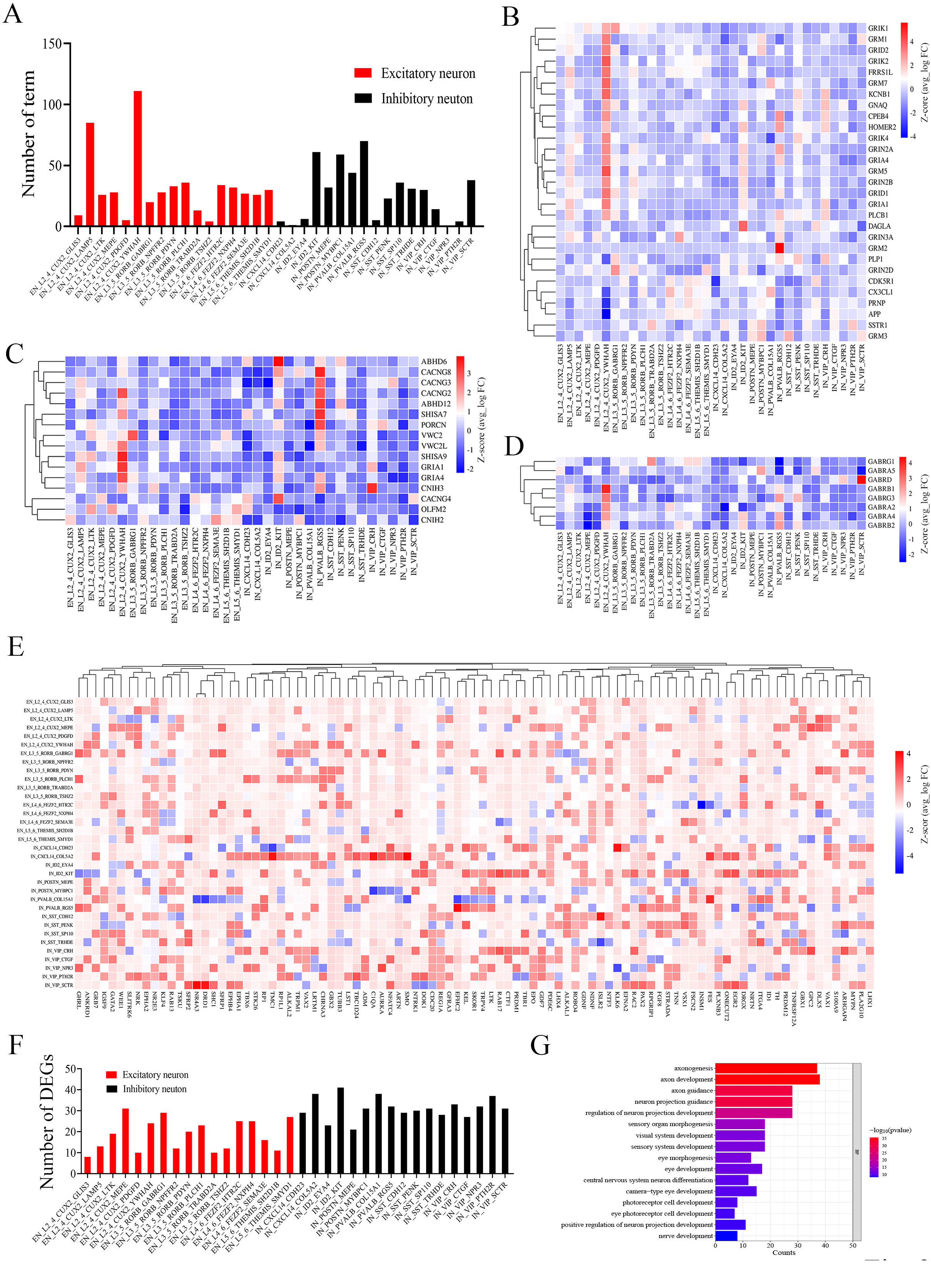
Major signaling pathways affected in FCD patient brain tissues. **(A)** Numbers of enriched neuron-related GO terms in neuronal subtypes. **(B)** Relative expression of DEGs in the glutamate receptor signaling pathway (Ep vs. Ctrl). **(C)** Relative expression of DEGs in the AMPA glutamate receptor complex (Ep vs. Ctrl). **(D)** Relative expression of DEGs in the GABA receptor complex (Ep vs. Ctrl). **(E)** Relative expression of DEGs in the neuron development pathways (Ep vs. Ctrl). **(F)** The number of DEGs with high difference in expression level (avg_log_2_ (FC) > 16 or < −16, *p* < 0.05) in neuron development pathway. **(G)** GO functional analysis of DEGs with higher difference in expression level. For B, C, D and E, colors represent the scaled average of Z-score (avg_log_2_ (Fold-Change). Fold-Change = G1/G2, with G1 and G2 represent the gene expression levels in epilepsy and control groups, respectively.

We have also examined signaling pathways associated with excitatory/inhibitory (E/I) balance.^47^ For glutamate receptor signaling pathway (GO:0007215), AMPAR complex (GO:0032281) and GABA receptor complex (GO:1902710), significantly higher scores in epilepsy-subtypes in epilepsy patients were observed (Supplementary Fig. 4A, 4B, 4C). High expression of DEGs (Ep vs. Ctrl) for both glutamate receptor signaling and AMPAR complex pathway were found in most epilepsy-subtypes (Fig. 3B, 3C). For GABA receptor complex pathway, higher expression in the epilepsy-subtypes was seen in PVALB_RGS5 and L2_4_CUX2_YWHAH subtypes (Fig. 3D).

**Figure 4.**
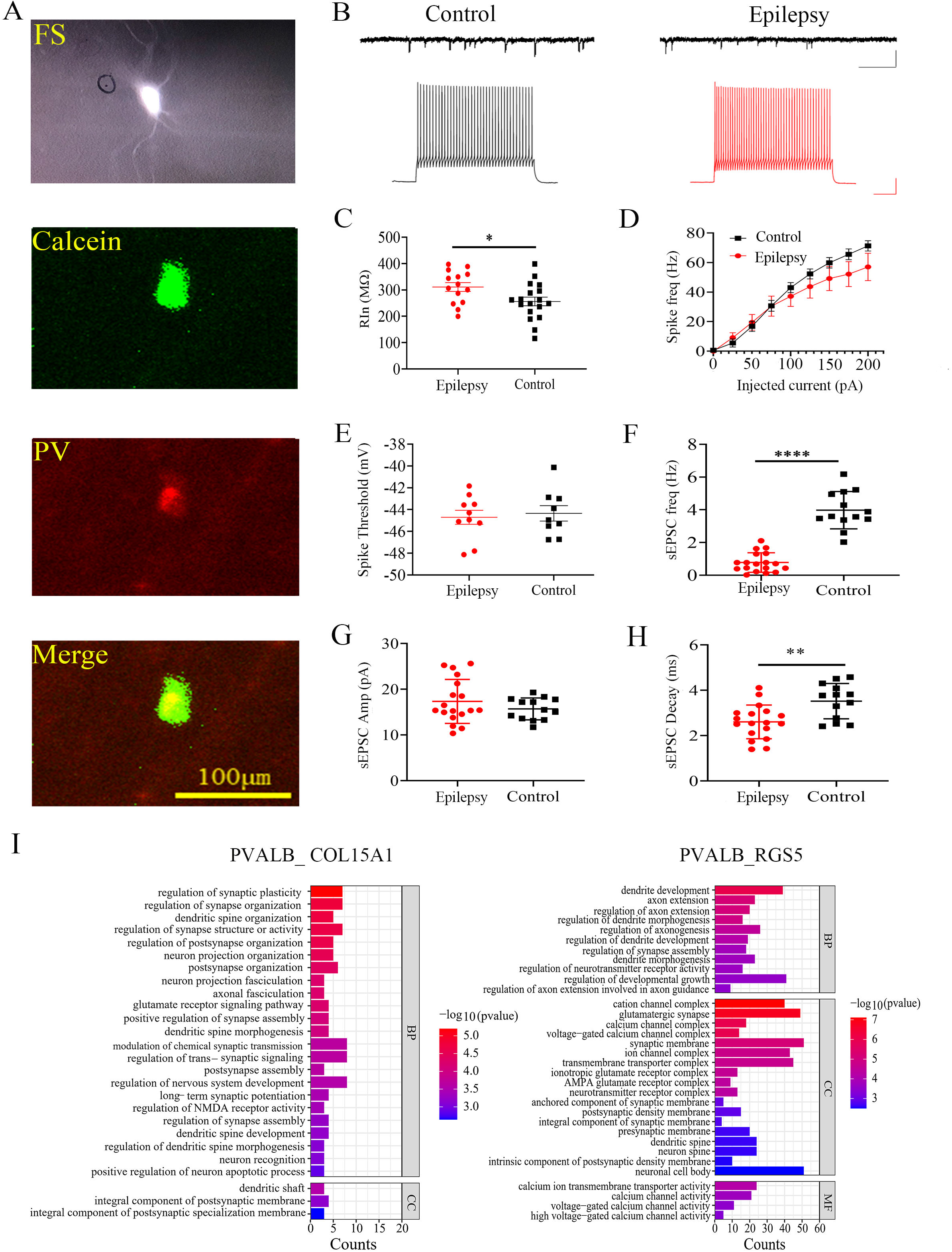
Alterations in the electrophysiological properties of fast-spiking neurons in child FCD patients. **(A)** Sample images of recorded fast-spiking neurons and immunostaining with anti-parvalbumin antibody after fixation. A green fluorescent dye (calcein) was included in the recording electrode, while red fluorescence indicates PV staining. Scale bar, 100 µm. **(B)** Sample traces of sEPSCs and spiking in FS-neurons in control or FCD groups. Scale bars, 50 pA/500 ms (sEPSCs) and 25 mV/100 ms (spikes). **(C)** Quantification of the input resistance of the recorded FS-neurons in epilepsy or control groups. **(D)** Spike frequencies to injection of depolarizing currents in FS-neurons in epilepsy or control groups. **(E)** Spike threshold in FS-neurons from epilepsy or control groups. **(F)** Frequency of sEPSC in FS-neurons from epilepsy or control groups. **(G)** The amplitude of sEPSC in FS-neurons from epilepsy or control groups. **(H)** The decay time of sEPSC in epilepsy and control groups. **(I)** GO enrichment analysis of DEGs in PVALB_COL15A1 and PVALB_RGS5 subtypes.

Analyzing the significant alterations in the aforementioned signaling pathways can identify dysregulated novel genes in epilepsy.^27^ We identified *GRIN3A*, *GRM5, GRIA1, GRID2, GRID1, GRIK4, GRM2, CDK5R1, DAGLA,* with the significant changes (avg_log_2_ (FC) > 1 or < −1, *p* < 0.05) in their expression in certain neuronal subtypes in glutamate receptor signaling pathway (Supplementary Table 5) and these genes had not been reported in pediatric FCD epilepsy patients. Both *GRIN3A* and *GRM2* were significantly up-regulated in L2_4_CUX2_YWHAH and PVALB_RGS5, while *GRM 5, GRIK4, GRID1* and *GRIA1* showed selective and significant up-regulation in L2_4_CUX2_YWHAH. For AMPA glutamate receptor complex pathway, we found *CACNG4, CNIH2, OLFM2, CACNG8, ABHD6, PORCN, SHISA7, VWC2, VWC2L, CNIH3, GRIA1,* and *SHISA*9 were associated with epilepsy in FCD patient tissues (Fig. 3C, Supplementary Table 6), and not reported in pediatric FCD patients previously. Many of these genes encode AMPA auxiliary subunits (*CACNG4, CNIH2, CACNG8, ABHD6, SHISA7, SHISA9*). *CACNG8,* and *SHISA7* were selectively and highly expressed in PVALB_RGS5, while *GRIA1* and *SHISA9* showed significantly higher expression in L2_4_CUX2_YWHAH subtype. For GABAR complex signaling pathway (Supplementary Table 7), only eight genes showed differential expression between epilepsy and control groups, with *GABRG3* and *GABRG1* under-studied in epilepsy.^48^ *GABRG3* was significantly up-regulated in the L2_4_CUX2_YWHAH, PVALB_RGS5, and SST_PENK subtypes. The L2_4_CUX2_YWHAH and PVALB_RGS5 subtypes exhibited the highest number of significantly altered genes (with an avg _log_2_ (FC) > 1 or < −1 and *p* < 0.05) in the above three receptor pathways, and suggest that these two subtypes have the most pronounced alterations in gene expression in the FCD patient brain.

We performed additional analysis on the neuronal development signaling pathways (GO:0048666) in the identified 34 neuronal subtypes. Unlike the gene set used in Figure 2C, the relevance of the majority of genes in the gene set used here to epilepsy has not been examined in patient population. We found a total of 1012 neuronal development-related genes showing altered expression in FCD patients (Supplementary Table 8). For the genes displaying the most significant differences in expression between the epilepsy and control groups (with an avg_log_2_ (FC) > 16 or < −16 and *p* < 0.05, Supplementary Table 9), particularly within the GABAergic subtypes (Fig. 3E, 3F). GO functional analysis revealed significant enrichment in axon-related, neuron projection and neuron development signaling pathways (Fig. 3G). Importantly, we found no relevance between the expression level of these identified genes and neuronal subtypes, indicating that these pan-neural development genes affect all neuronal subtypes non-selectively.

We also found DEGs (Ep vs. Ctrl) were enriched in neurodegeneration-multiple disease pathways in certain neuronal subtypes (Supplementary Fig. 4D). Certain pathways were unique for specific neuronal subtypes. For example, long-term potentiation pathway was selectively and significantly enriched in L2_4_CUX2_YWHAH, and HIF-1 signaling pathway was the only enriched in L3_5_RORB_GABRG1 subtype (Supplementary Fig. 4D). Furthermore, many of the same pathways were enriched in both L2_4_CUX2_YWHAH and PVALB_RGS5, including oxytocin signaling, circadian entrainment, dopaminergic synapse, and cholinergic synapse pathways, and supporting that these two subtypes are more likely affected in epilepsy. We also found certain addiction pathways, including morphine, nicotine and cocaine pathways, were enriched by DEGs (Ep vs. ctrl) in a several subtypes (Supplementary Fig. 4D). Put together, our analysis showed that DEGs in epilepsy-related subtypes were enriched in neuronal function-related and E/I balanced-related pathways. We have discovered several novel genes that are specific to certain subtypes and are associated with E/I balance pathways. These genes have not been previously reported in pediatric drug-resistant FCD epilepsy. Our results also support that larger numbers of epileptogenesis - related genes are enriched in both L2_4_CUX2_YWHAH and PVALB_RGS5 subtypes.

### Alterations in fast spiking/parvalbumin-neurons in FCD brains

Findings from Figure 2 and Figure 3 indicate that PVALB_RGS5 is one of the most prominent epilepsy-subtypes. The PV-neurons are the most widely studied neuronal subtype in epilepsy,^49,50^ as their potent inhibition can efficiently suppress neuronal spiking and prevent seizure onset.^49^ A high percentage of PV-neurons display high frequency spiking (fast-spiking, FS) to injection of depolarizing current, and hence they can be identified using patch-clamp recording in acute brain slices. We recorded from FS-neurons in acute brain slices from both epilepsy and control zone (see Methods) (Fig. 4, Supplementary Fig. 5). We included a green fluorescent dye in the recording solution which diffused into the recorded neurons during the recording session. We then performed staining using PV-selective antibodies (red fluorescence) on the fixed slices. Overlap of green and red fluorescence within a single neuron confirmed the recorded neuron as PV-positive (Fig. 4A). We found no difference in the intrinsic excitability of FS-neurons between epilepsy and control groups (Fig. 4B, 4D). A significantly higher input resistance (Fig. 4C), a lower frequency and decay time of sEPSC (Fig. 4F, 4H) was found in the epilepsy group, but these was no difference in spike threshold (Fig. 4E) or sEPSC amplitude (Fig. 4G). These results indicate that certain intrinsic neuronal and synaptic properties are alerted in the FS/PV-neurons in the FCD patient brains.

Since only one of the two PV-neuron subtypes is identified as epilepsy-subtype, we further examined transcriptomic differences between these two PV-subtypes in epilepsy. For PVALB_ COL15A1, DEGs between epilepsy and control groups were mainly enriched in pathways related to synaptic structure and glutamate receptor signaling (Fig. 4I, Supplementary Fig. 6A), while DEGs were enriched in pathways related to ion channels, axon structure, dendrite function, and AMPA receptor complex for PVALB_RGS5 (Fig. 4I, Supplementary Fig. 6B). Altogether, genes associated with glutamate receptor and GABA receptor signaling pathways were altered in the PV-neurons and excitatory synaptic inputs were reduced in these neurons. These alterations implicate reduced activity in these neurons and likely higher E/I ratio in epilepsy brains.

### Altered ATP production and energy metabolism in FCD brains

Impaired bioenergetics in epilepsy manifests to breakdown of mitochondrial machinery.^51^ We found that the averaged percentage of mitochondria gene expression level was higher in the epilepsy-subtypes, especially in L2_4_CUX2_YWHAH, ID2_KIT, PVALB_RGS5 or VIP_CRH subtypes (Fig. 5A). Importantly, this finding holds true for most individuals (Supplementary Fig. 7 and 8), suggesting that they are common. Furthermore, by using U Cell Gene set scoring method, we found a higher score in both oxidative phosphorylation signaling (Supplementary Fig. 9A) and ATP biosynthesis pathway (Supplementary Fig. 9C) in most epilepsy-subtypes. We speculate that neuronal subtypes displaying higher averaged percentage of mitochondria gene expression levels are more vulnerable to epilepsy genesis. However, comparing between epilepsy and control groups, we found a significantly lower scores in both oxidative phosphorylation (has00190) (Fig. 5B, S9B) and ATP biosynthesis (GO:0006754) signaling pathways (Fig. 5C, Supplementary Fig. 9D) in most epilepsy-subtypes from epilepsy groups, especially in L2_4_CUX2_YWHAH (score ratio: −12 and −15) and PVALB_RGS5 subtypes (score ratio: −5 and −18) (Fig. 5D, 5E). This finding suggests an impaired ATP production in these two epilepsy-subtypes in epilepsy patient brains. For the other epilepsy-subtypes, the impairment in ATP production pathway occurred to a less extent or not at all. This subtype selectivity may explain the differences in ATP level during seizure in previous studies.^52^ Taken together, these observations indicate that in normal brains, ATP/mitochondria functions are more pronounced in the epilepsy-subtypes compared to other subtypes. Furthermore, the lower activity of ATP pathway in the epilepsy-subtypes in the FCD brains suggests that a link between seizure and impaired ATP production in these subtypes. This lower function in ATP pathway predicts a reduced metabolic activity in the epilepsy brain regions, which is supported by the hypometabolism in the epileptogenic zone in our pediatric FCD patients (Supplementary Table 1) and in 70 - 89% of patients with drug-resistant epilepsy,^53^ measured using [^18^F]-fluorodeoxyglucose (FDG) positron emission tomography.

**Figure 5.**
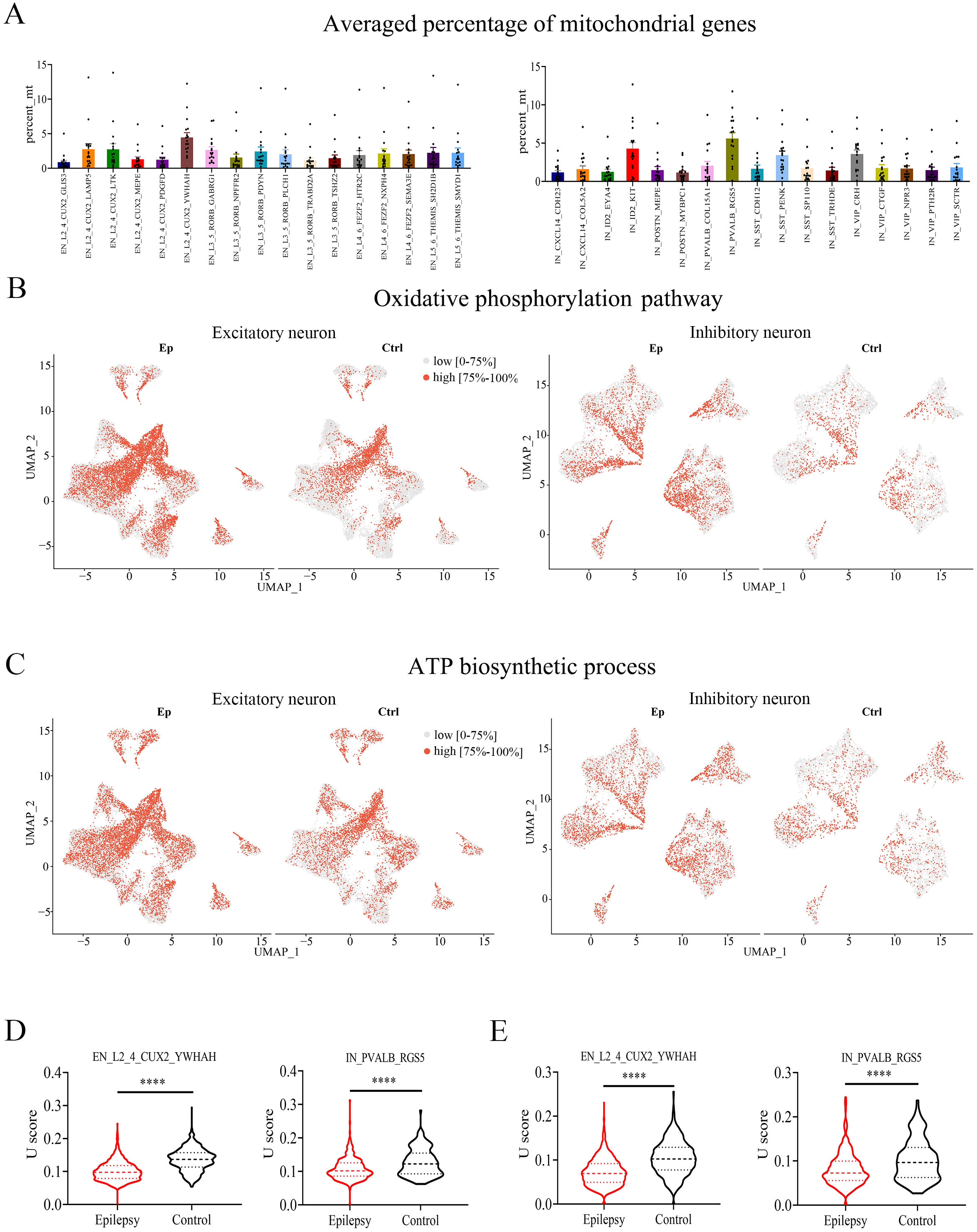
Genes associated with mitochondrial functions are altered in FCD epilepsy patients. **(A)** Averaged percentage of mitochondrial genes expression in excitatory neuron and inhibitory neuronal subtypes. **(B)** U scores of oxidative phosphorylation pathway in the excitatory and inhibitory neuronal subtypes between epilepsy and control groups. **(C)** U scores of ATP biosynthetic process in the excitatory and inhibitory neuronal subtypes between epilepsy and control groups. **(D)** U scores of oxidative phosphorylation in the L2_4_CUX2_YWHAH and PVALB_RGS5 subtypes between epilepsy and control groups. **(E)** U scores of ATP biosynthetic process in the L2_4_CUX2_YWHAH and PVALB_RGS5 subtypes between epilepsy and control groups. For B and C, red represents high scores (top 75%) and gray represents low sores (0 - 75%).

## Discussion

By using snRNA-seq, we found that out of the 17 excitatory and 17 inhibitory neuron subtypes, only a few displayed significant epilepsy-related transcriptomic alterations, in the pediatric drug-resistant FCD epilepsy patients. In addition, DEGs associated with neuronal functions and E/I balance exhibit the most prominent alterations in these specific subtypes. Supporting PVALB_RGS5 PV-neurons being a prominent epilepsy-subtype, we observed lower excitatory synaptic inputs in PV-neurons in acute FCD patient brain slices. Furthermore, a higher percentage of genes related to mitochondria and ATP function were altered in the most epilepsy-subtypes, especially in L2_4_CUX2_YWHAH and PVALB_RGS5 subtypes.

Our findings support a prior report that selective neuronal subtypes have preferentially higher contributions to epilepsy genesis in the adult epilepsy patients.^27^ However, the subtypes they have identified differ from ours, which may be caused by differences in patient age, detailed pathology and analysis method used. We identified two PV-neuron subtypes, with only PVALB_RGS5 subtype met the criteria for epilepsy-related subtype, and this conclusion is supported by the larger number of epilepsy-related genes exhibiting higher altered expression and the presence higher number of DEGs within the E/I-related pathways in this particular subtype. Altered expression of both glutamate receptors and AMPAR auxiliary subunits was also observed in this PV-subtype, consistent with an altered/impaired glutamatergic transmission onto PV-neurons that we have observed in FCD patient brain slices. Since these samples were obtained from pediatric patients, altered development/maturation of inhibitory neurons may constitute a major cause for epilepsy in pediatric FCD patients. It will be interesting and important to understand why only one of the two PV-neuron subtypes is epilepsy-related subtype, and whether selective targeting this subtype may provide effective treatment for FCD.

Another very interesting and potentially important findings is a higher average percentage of mitochondria genes in the epilepsy-subtypes. Mitochondrial dysfunction has been proposed as a contributor to seizure initiation or a consequence of excessively prolonged seizures.^51,54,55^ During epilepsy, prolonged repetitive neuronal activity aggravates the demand of energy use and disrupts the balance in bioenergetics leading to energy crisis and ATP shortage.^51,56,57^ Mitochondrial dysfunction may also lead to reduced energy supply and subsequent neuronal dysfunction. The higher average percentage of mitochondria gene expression in the epilepsy-subtypes in both control and epilepsy brains suggest that these subtypes may require higher energy to maintain their normal functions, and shortage in energy may render them more vulnerable to epilepsy.

Lower scores in oxidative phosphorylation and ATP biosynthesis pathways in the PVALB_RGS5 subtype in FCD patients suggests that the high demand for mitochondrial energy metabolism in the PV-neurons may not be adequately met in the epilepsy brains.^20^ Further examination is required to determine whether altered ATP biosynthesis pathways contribute to the cause, consequence, or compensation processes in the epilepsy.^23,24,52,58^ Supporting the critical contribution of mitochondria genes in epilepsy, we found higher scores in both oxidative phosphorylation and ATP biosynthesis pathways in most epilepsy-subtypes when comparing to the other subtypes, while scores in the epilepsy-subtypes were lower in the epileptic tissue when comparing to non-epileptic tissue. Why are only a handful neuronal subtypes displaying high association with epilepsy? A few possibilities are worthy of consideration: (1) the higher energy demands or lower mitochondria/ATP functions in these epilepsy-subtypes may render them more vulnerable to the increase in activity or energy use in the brain. The altered E/I-related pathways in these epilepsy subtypes likely lead to higher brain excitation, which in turn induces higher brain activity and energy demand. Alternatively, epilepsy may occur as a consequence of impaired energy adaptation, particularly in the inhibitory neurons. This impairment can lead to insufficient inhibition, thereby compromising the ability to control seizures effectively. (2) The PVALB_RGS5 subtype may exert more powerful and effective control over brain excitation and preventing the runaway excitation, in comparison to the other PV-subtype and other inhibitory neuron subtypes. This suggests that manipulating the PVALB_RGS5 -subtype may be more effective in seizure prevention or control. (3) These epilepsy-subtypes are developmentally more vulnerable to alterations or perturbations in the environment, especially during the protracted cortical development. This scenario may be particularly relevant for the inhibitory neurons. It is important to note that the transcriptomic findings we reported here alone cannot determine the sequence or causal relationship of the implicated processes. Perturbation experiments are required to investigate this question. Furthermore, it is possible that certain alterations may be common in the majority of FCD patients, but the initial causes may differ.

In conclusion, our snRNA-seq analysis on freshly obtained pediatric drug-resistant FCD epilepsy patients reveal prominent contributions in only a few selective neuronal subtypes. In these epilepsy-subtypes, the genes associated with a few selective functions/signaling pathways display the most prominent alterations. Among the identified genes/pathways, further investigation is necessary to explore the involvement of mitochondria and ATP signaling pathways in FCD epilepsy. These new findings offer a foundation for gaining deeper insights into the classification, specificity, and underlying disease mechanisms of epilepsy, as well as potential improvement of diagnostics and more effective treatments for epilepsy.

## Acknowledgements

We thank all the patients and clinicians that participated in this work.

## Supplementary material

Supplementary material is available online.

## Supplementary materials

**Supplementary Fig 1.**
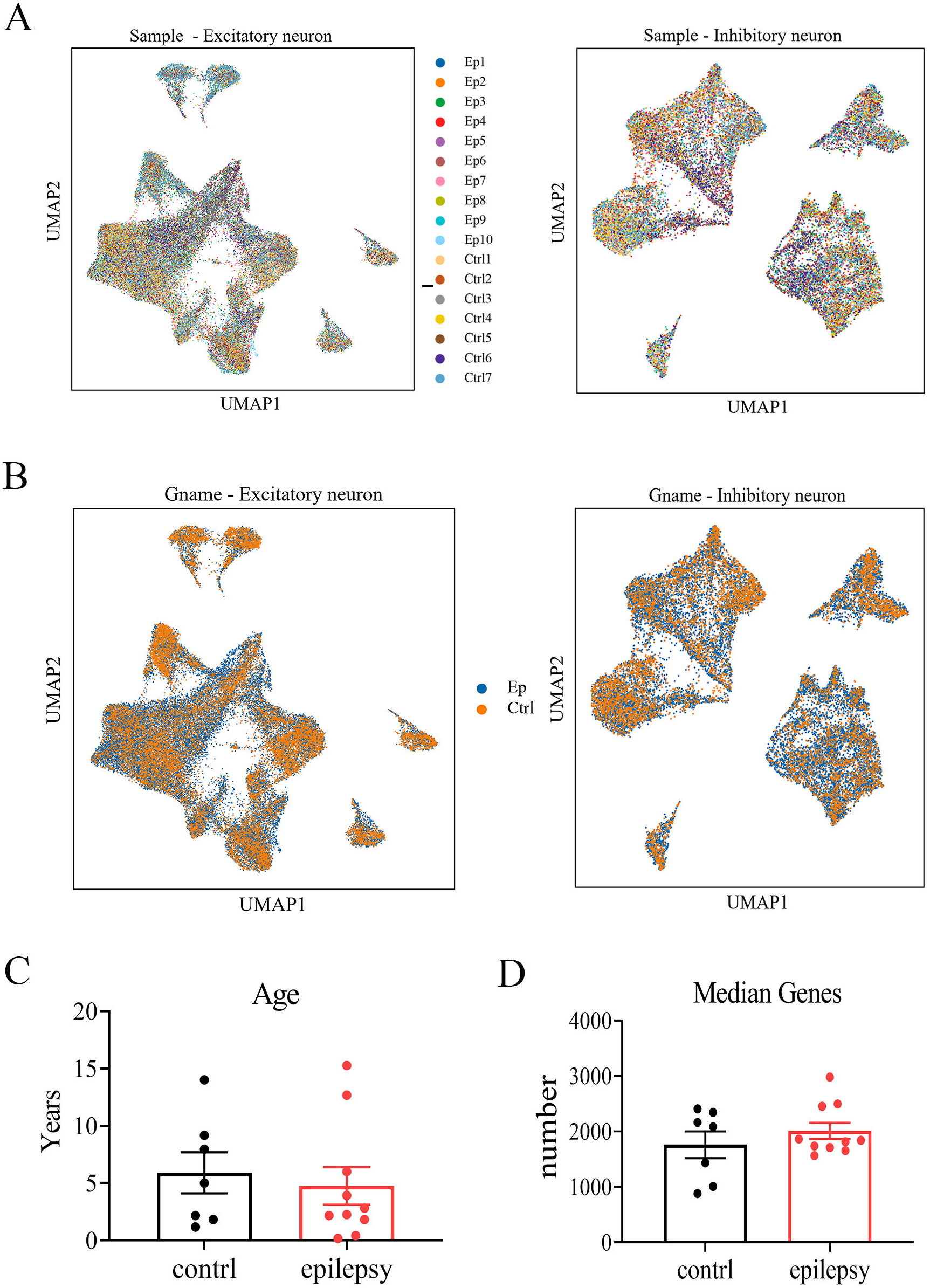
Information on epilepsy and control patient groups. **(A)** UMAP representation of excitatory and inhibitory neuronal in all patient samples. **(B)** UMAP representation of excitatory and inhibitory neuronal between epilepsy and control groups. **(C)** Age of all FCD patients. **(D)** Median gene numbers from all patients.

**Supplementary Fig 2.**
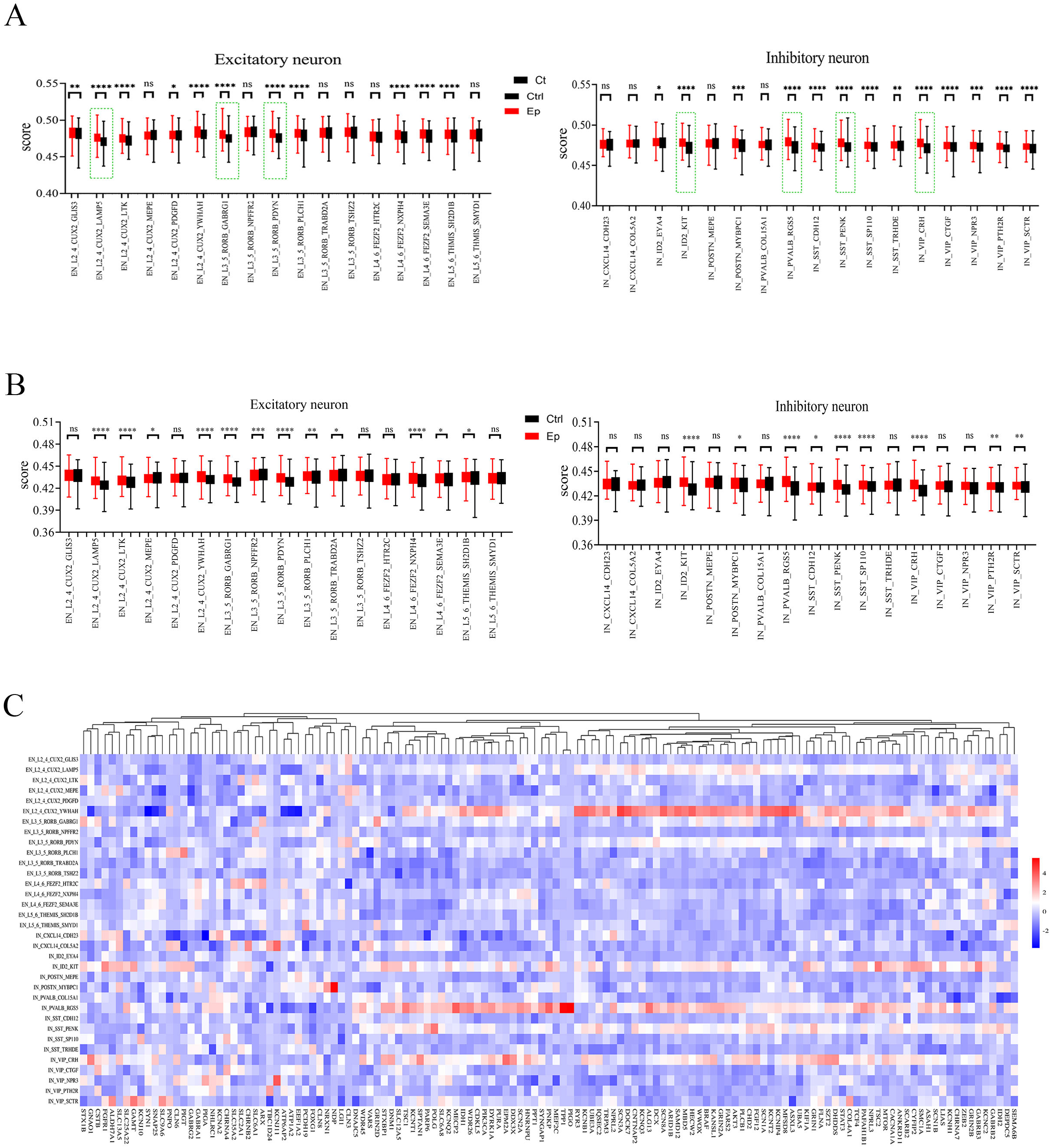
Expression of epilepsy-associated genes in neuronal subtypes. **(A)** U scores of epilepsy genes identified in DisGeNET (C10012544) in excitatory neurons and inhibitory neurons between epilepsy and control groups. The green dotted line was the neuronal subtypes with a high score ratio. **(B)** U scores of epilepsy-related genes in excitatory and inhibitory neuronal subtypes between epilepsy and control groups. **(C)** Relative expression levels (Ep vs. Ctrl) of 143 epilepsy-associated genes in excitatory or inhibitory neuronal subtypes.

**Supplementary Fig 3.**
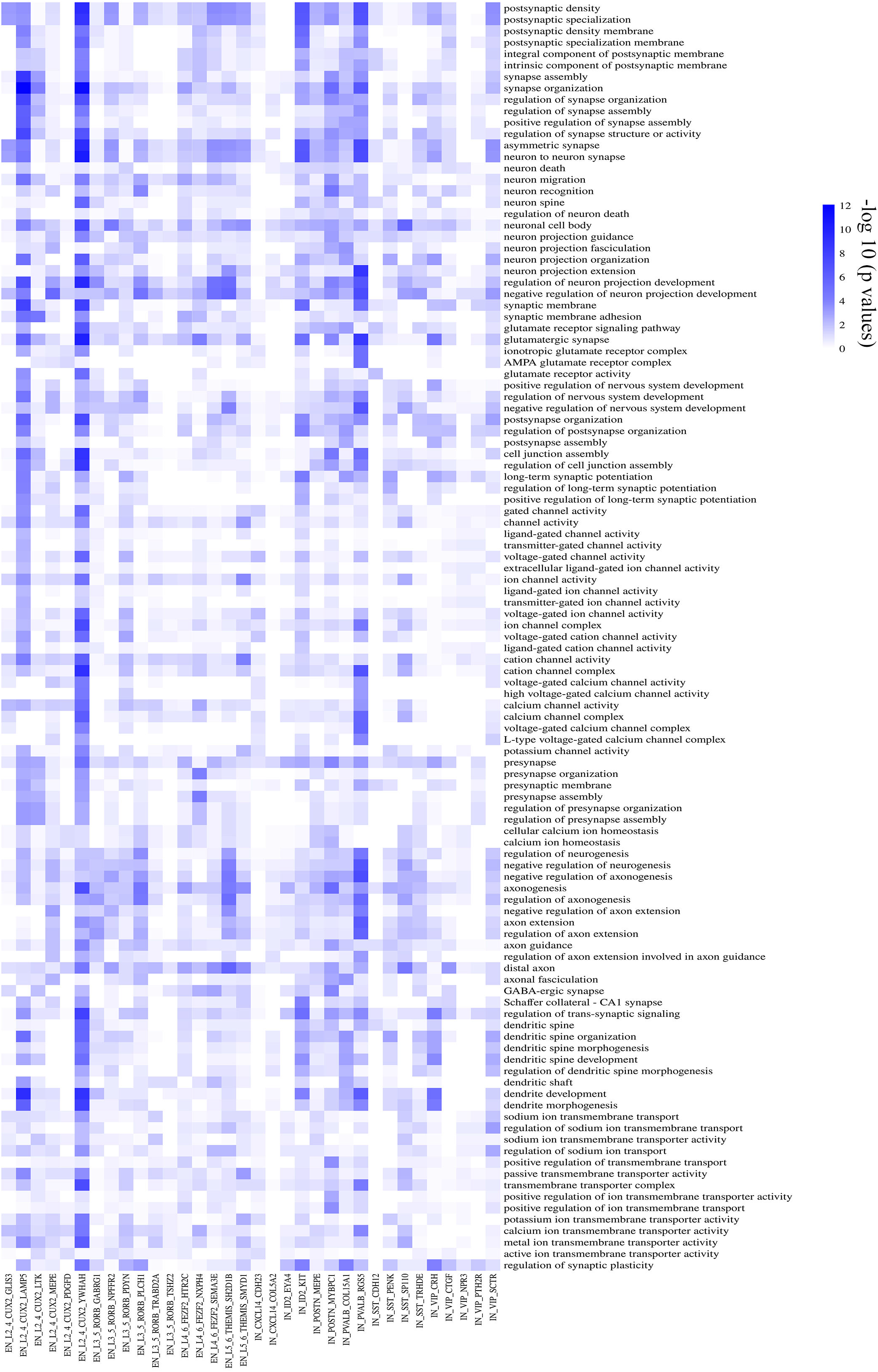
GO functional enrichment analysis of DEGs on neuronal function-related pathways. Colors represent –log_10_ (*P* values).

**Supplementary Fig 4.**
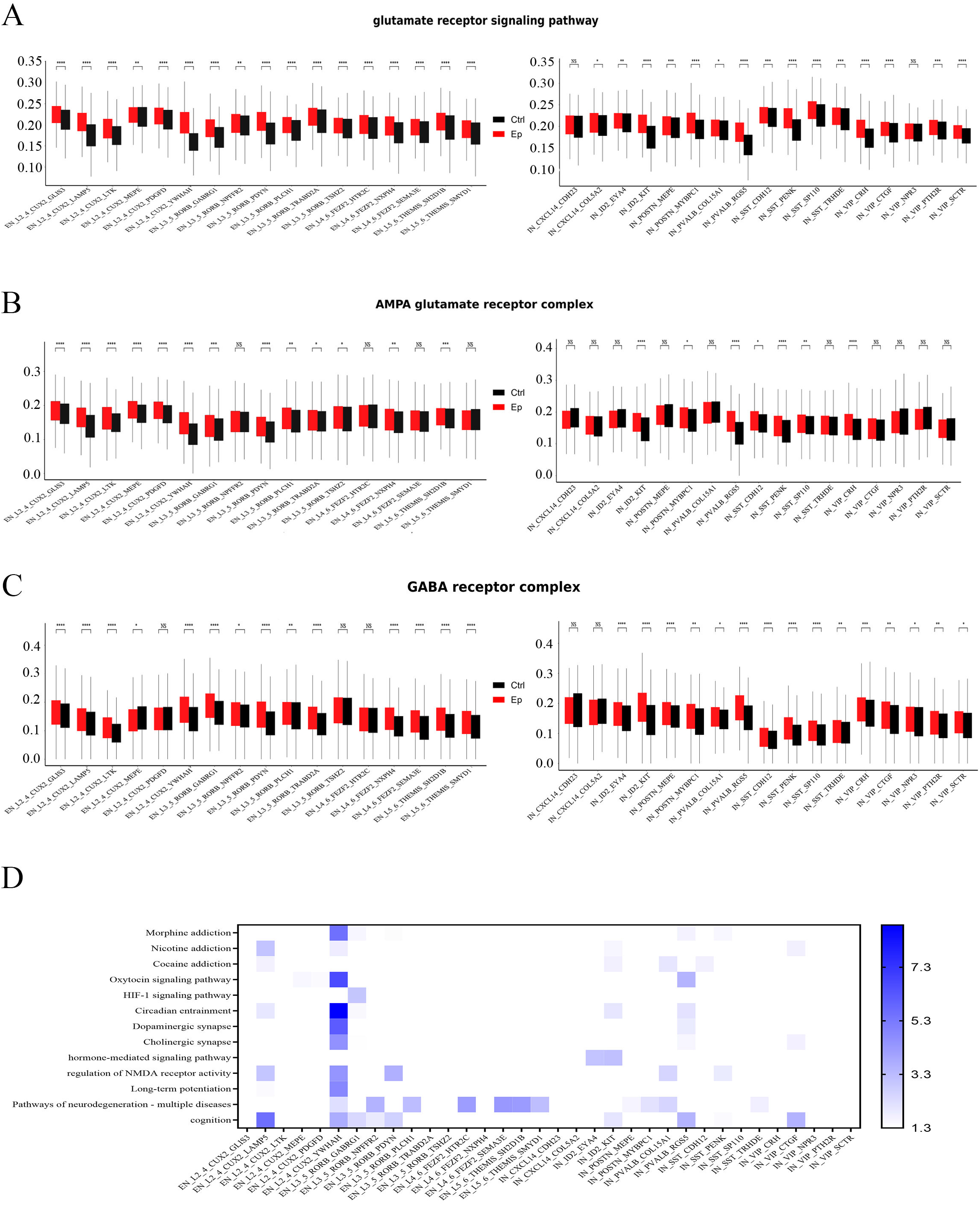
The activity of selective signaling pathways in neuronal subtypes. **(A)** U scores of glutamate receptor signaling pathway in the excitatory and inhibitory neuronal subtypes between epilepsy and control groups. **(B)** U scores of AMPAR complex pathway in the excitatory and inhibitory neuronal subtypes. **(C)** U scores of GABA receptor complex pathway in the excitatory and inhibitory neuronal subtypes. **(D)** Selective pathways enriched in certain neuronal subtypes.

**Supplementary Fig 5.**
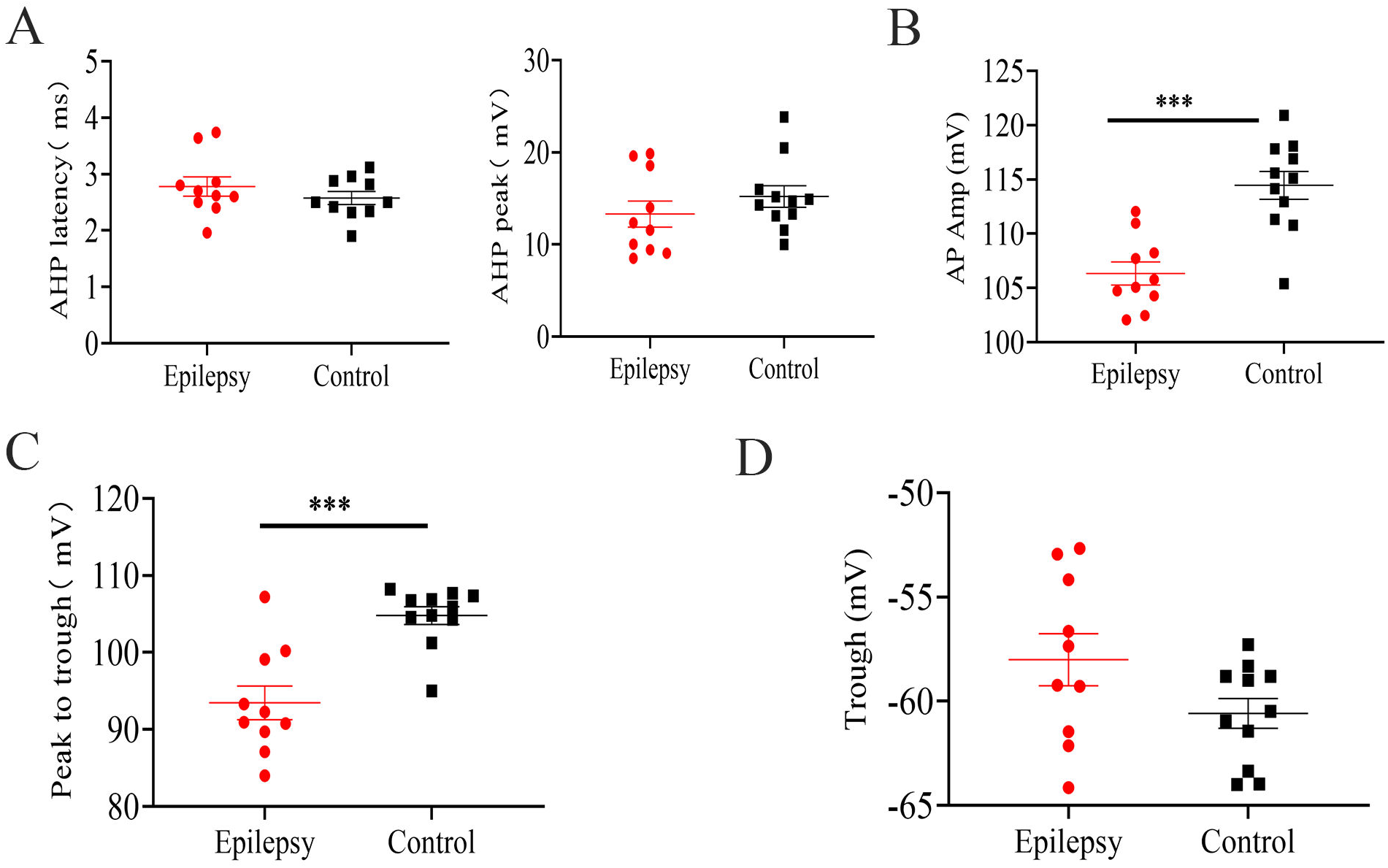
Electrophysiology properties of FS-neurons. **(A)** AHP latency (left) and peak (right) in FS-neurons from epilepsy and control groups. **(B)** Quantification of spike amplitude in FS-neurons from epilepsy and control groups. **(C)** Quantification of peak to trough amplitude in spikes of FS-neurons in epilepsy and control groups. **(D)** Quantification of spike trough amplitude in FS-neurons from epilepsy and control groups.

**Supplementary Fig 6.**
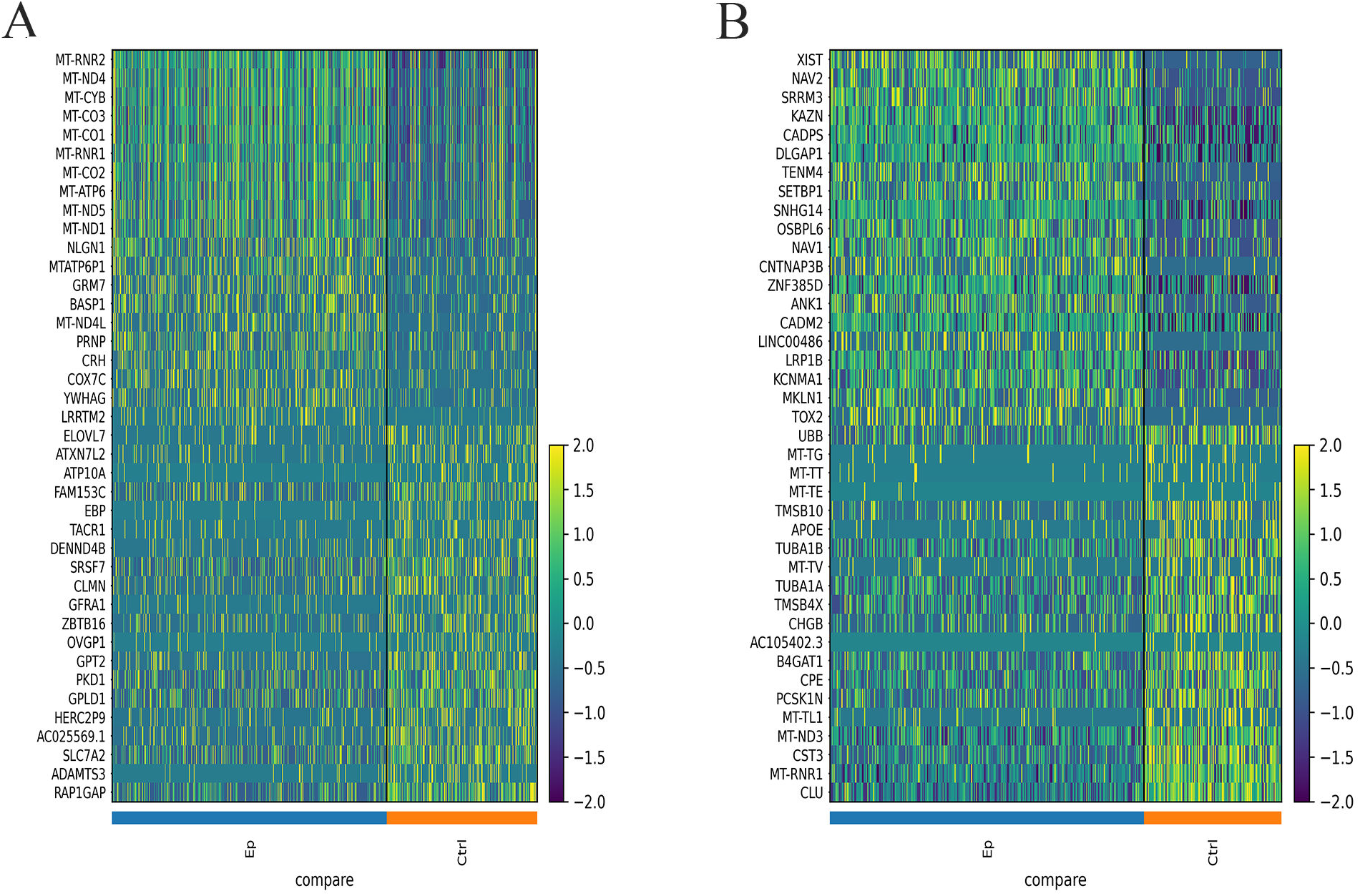
The top 20 DEGs of both two PV subtypes between epilepsy and control group. **(A)** Top 20 up- or down-regulated DEGs in the PVALB_COL15A1 subtype that were significantly different between epilepsy and control groups. **(B)** Top 20 up- or down-regulated DEGs in the PVALB_RGS5 subtype that were significantly different between epilepsy and control groups.

**Supplementary Fig 7.**
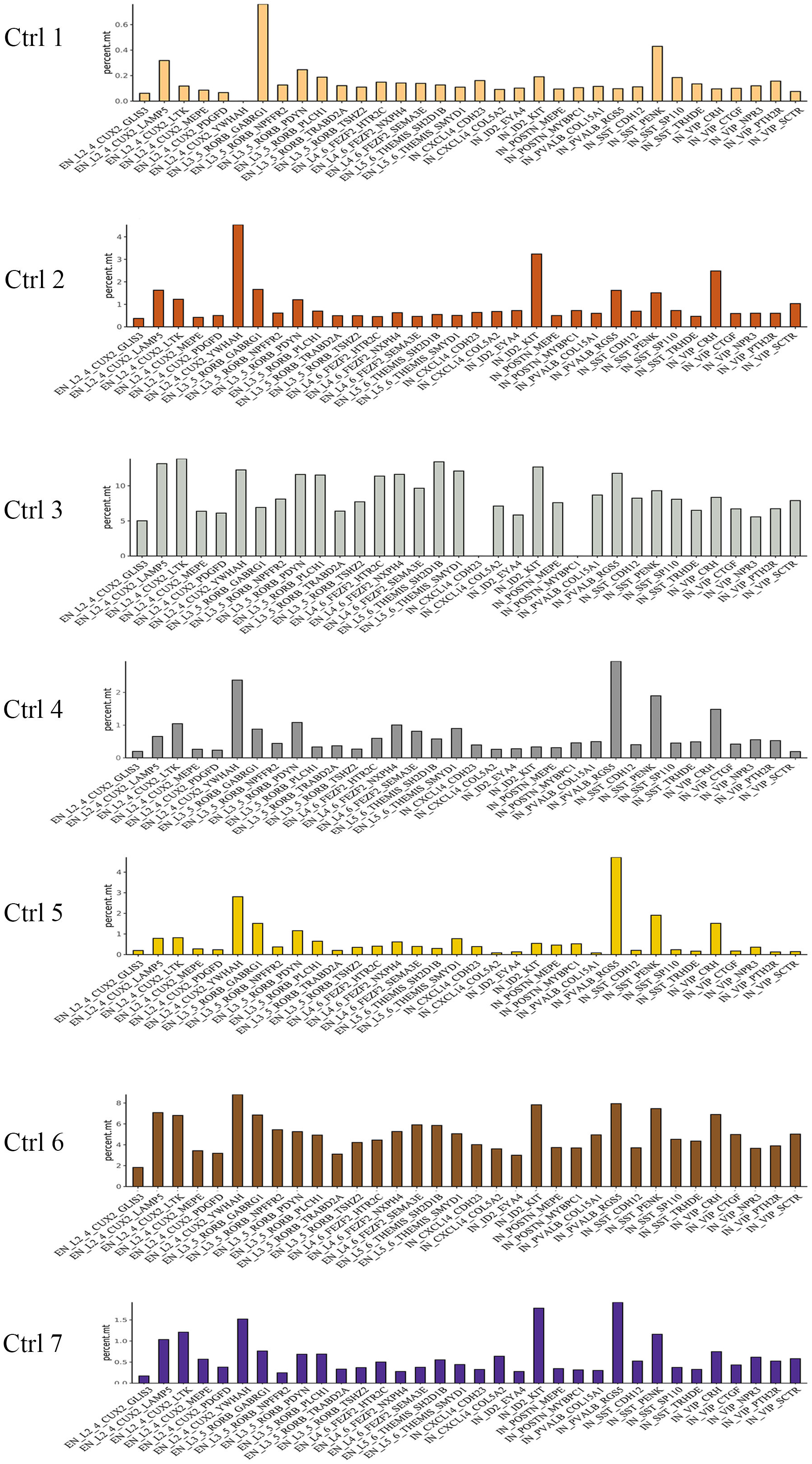
Averaged percentage of mitochondrial genes in neuronal subtypes in individuals of the control group.

**Supplementary Fig 8.**
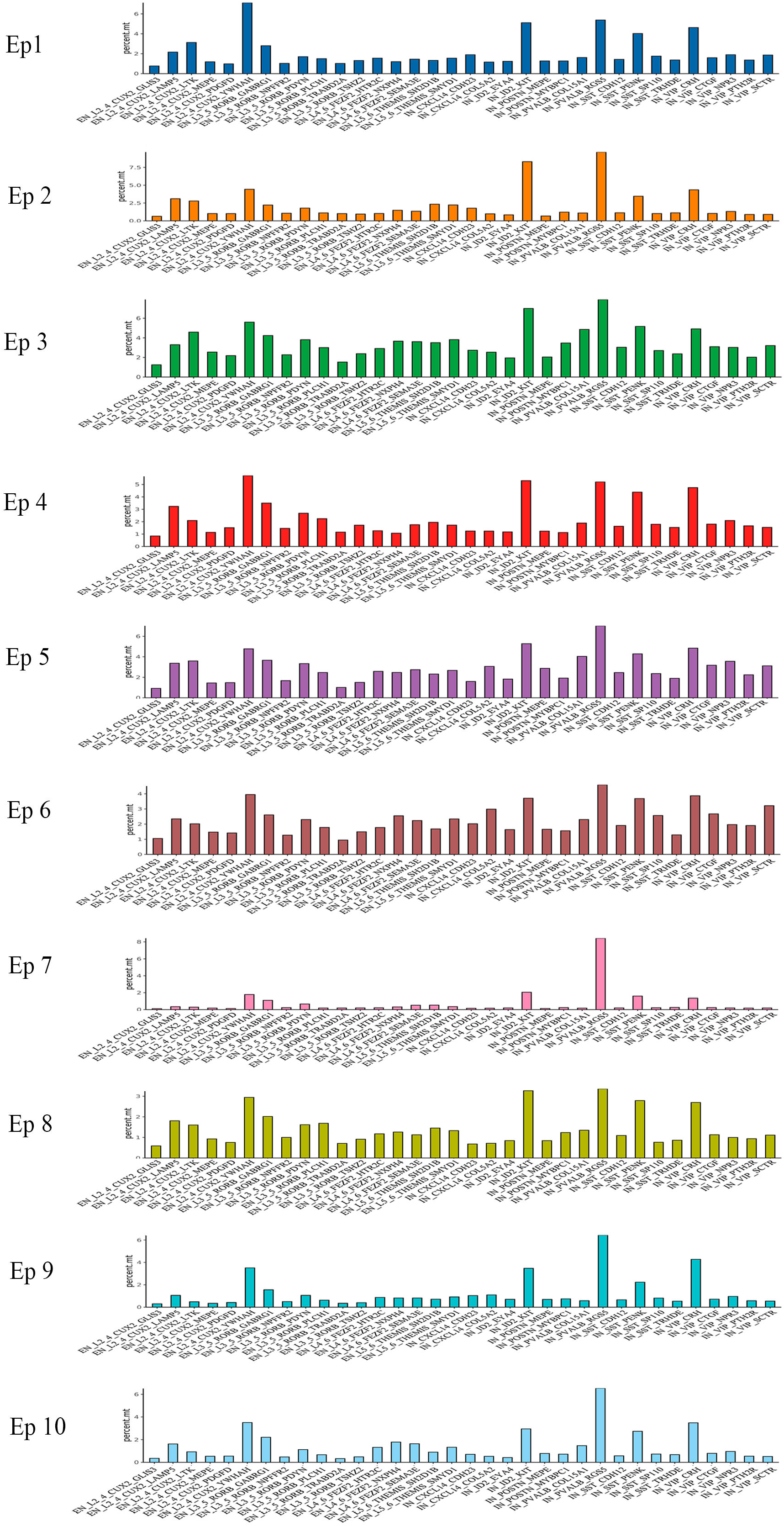
Averaged percentage of mitochondrial genes in neuronal subtypes in individuals of the epilepsy group.

**Supplementary Fig 9.**
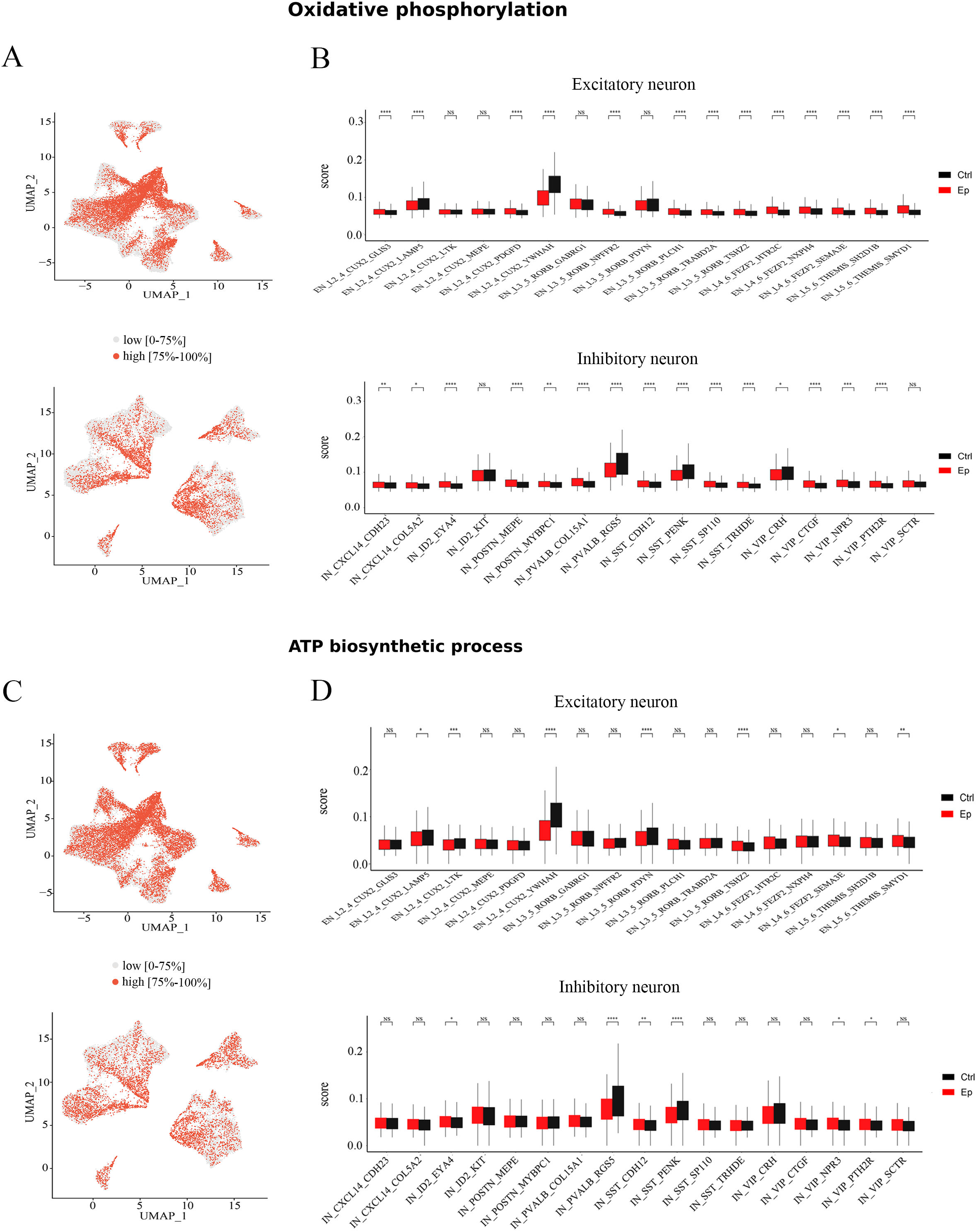
The activity of oxidative phosphorylation and ATP biosynthetic process in neuronal subtypes. **(A)** UMAP representation of the sore of oxidative phosphorylation pathway in neuronal subtypes. Red represents high scores (top 75%) and gray represents low sores (0 - 75%). **(B)** U scores of oxidative phosphorylation pathway in excitatory and inhibitory neuronal subtypes between epilepsy and control groups. **(C)** UMAP representation of the sore of ATP biosynthetic process pathway in neuronal subtypes. Red represents high scores (top 75%) and gray represents low sores (0 - 75%). **(D)** The U scores of ATP biosynthetic process pathway in excitatory and inhibitory neuronal subtypes between epilepsy and control groups.

